# Meta-Learning Biologically Plausible Semi-Supervised Update Rules

**DOI:** 10.1101/2019.12.30.891184

**Authors:** Keren Gu, Sam Greydanus, Luke Metz, Niru Maheswaranathan, Jascha Sohl-Dickstein

## Abstract

The question of how neurons embedded in a network update their synaptic weights to collectively achieve behavioral goals is a longstanding problem in systems neuroscience. Since Hebb’s hypothesis [10] that cells that fire together strengthen their connections, cellular studies [6] have shed light on potential synaptic mechanisms underlying learning. These mechanisms have directly driven the careful hand design of biologically plausible models of learning and memory in computational neuroscience [1]. However, these hand designed rules have yet to achieve satisfying success training large neural networks, and are dramatically outperformed by biologically implausible approaches such as backprop. We propose an alternative paradigm for designing biologically plausible learning rules: using meta-learning to *learn* a parametric synaptic update rule which is capable of training deep networks. We demonstrate this approach by meta-learning an update rule for semi-supervised tasks, where sparse labels are provided to a deep network but the majority of inputs are unlabeled. The meta-learned plasticity rule integrates bottom-up, top-down, and recurrent inputs to each neuron, and generates weight updates as the product of pre- and post- synaptic neuronal outputs. The way in which the inputs to each neuron are combined to produce a learning signal, however, is itself a *meta-learned* function, parameterized by a neural network. Critically, the meta-learned update rule integrates only neuron-local information when proposing updates–that is, our learning rule is spatially localized to individual neurons. After meta-learning, the resulting synaptic update rule is capable of driving task-relevant learning for semi-supervised tasks. We demonstrate this capability on two simple classification problems. In general, we believe meta-learning to be a powerful approach to finding more effective synaptic plasticity rules, which will motivate new hypotheses for biological neural networks, and new algorithms for artificial neural networks.

## 1 Introduction

Our approach is to directly optimize a synaptic weight update rule using a meta-objective that maximizes performance on a suite of semi-supervised learning tasks. Additionally, we design our learning rule to be biologically plausible: it is *neuron local*, meaning updates to weights depend only on the corresponding pre- and post- synaptic units; it further does not use weight tying between feedforward and feedback passes during training [16]. These design features enable the update rule to be applied to train models with different depths or widths, and to be applied to new datasets. In contrast to previous work [18], our approach is more biologically plausible, simpler, and tackles semi-supervised rather than unsupervised learning. We show that meta-learned biologically plausible learning rules for semi-supervised learning can outperform supervised learning on 2D toy tasks, and have the ability to generalize to unseen tasks.

## 2 Related Work

### 2.1 Learning local learning rules

Understanding how networks of interconnected neurons can learn to solve tasks using only information that is spatially and temporally local to individual neurons is an active area of research in both neuroscience and machine learning [23]. Prior research along these lines have proposed and studied specific local learning rules [3] or approximated existing non-local algorithms (such as backpropagation) with more biologically plausible variants [16, 25, 20].

Our work here follows a different thread, which uses meta-learning to learn a local learning rule. Learning a parameterization of a local synaptic learning rule was originally proposed in the early 1990s [5, 4]. Despite differences in scale (for example, these early learning rules had just 7 or 16 meta-parameters [4]), the overall setup is largely similar: write down a parameterized learning rule that is a function of neuron local variables, specify a set of training task(s), and finally optimize the learning rule for those tasks using either 1st (gradient descent) or 0th order optimization methods (simulated annealing, genetic algorithms). Concurrent work along these lines uses multilayered fully connected networks to parameterize the learning rule [17]. A distinct line of research does not focus on biological plausibility, that is, they use meta-learning for learning non-local learning algorithms [12, 21, 22, 2, 24].

### 2.2 Semi-supervised learning

Semi-supervised learning (SSL) refers to a class of approaches that use unlabeled data to improve supervised learning. One approach to SSL starts by learning an unsupervised representation using unlabeled data, followed by fine tuning the model on a small labeled dataset [7, 8, 11]. Another common approach to SSL uses unlabeled data during training by adding additional loss terms, for example via generative modeling [13, 14], entropy minimization [9], small perturbations [19], or creating labels for nearby unsupervised examples [15].

## 3 Methods

Our meta-training procedure consists of a learning loop (applying the learning rule to update the weights of a network), which itself is wrapped in a meta-learning loop (Fig. 1, Left). The training procedure involves iteratively improving the learning rule in the meta-learning loop, where each iteration involves training and evaluation of the learning rule on a sampled task. In a single iteration of meta-training, we sample data from a semi-supervised learning task (discussed below), initialize a base network with random weights, and perform a fixed number of learning iterations where the learning rule is applied to update the base network weights. The inner model (the model the learned update rule is training) is a 1-hidden layer fully connected neural network in all experiments, with feed-forward weights *W*. We introduce two additional sets of weights 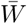 and 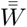 in addition to the feed forward weights *W* (Fig. 1, Right). These two sets of auxiliary weights help propagate error top-down in the network 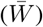 and communicate lateral signals between neurons in the same layer 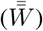, and are exclusively used for learning and not prediction. These are loosely inspired by the numerous feedback and recurrent connections present in cortex. All three sets of weights are initialized at random and updated using the learned update rule.

**Figure 1:**
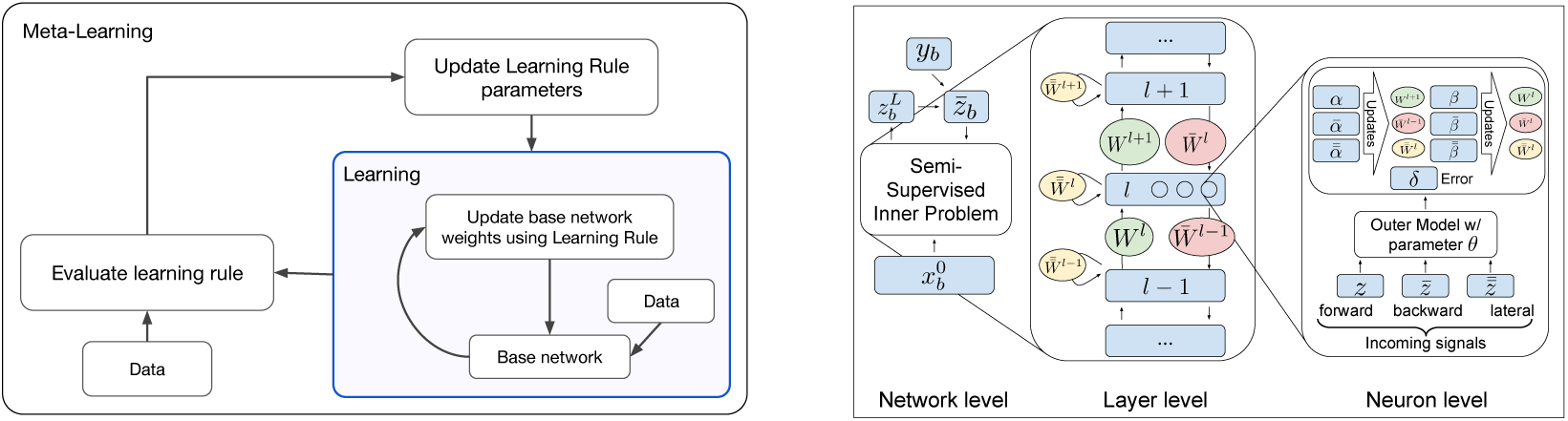
**Left:** Schematic of inner and outer learning loops in meta-training. **Right:** Schematic showing how the learned neuron-local semi-supervised learning rule produces weight updates.

Instead of computing the gradient of the parameters with respect to the loss (as would be done in backprop), we query the learned update rule for updates to the inner-parameters. Specifically, we compute updates one layer at a time from the top of the network down. For each neuron *i*, we use the forward activation (*z*), reverse signal (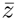 obtained from 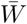), and lateral signal (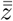 obtained from 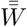) to query the outer model and compute weight updates 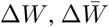, and 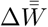 for pre- and post- synaptic neurons *j* and *i*:

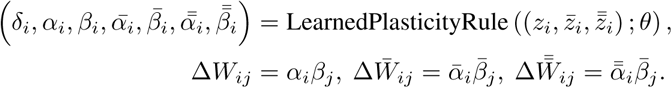

The output *δ* from the outer model is a learned top down error signal, and *θ* denotes the meta-parameters that parameterize the learning rule. See the Appendix A for a detailed description for how weight updates, and reverse and lateral signals, are computed.

## 4 Results

To generate our preliminary results, we train a fully connected neural network with one hidden layer. We train and evaluate semi-supervised learning rules on toy classification tasks. We train on tasks formed by re-sampling labeled examples from the two-moons dataset.

For the inner loop, we use a batch size of 10 and train for 50 steps. We compare supervised learning from 10 labeled examples (using backprop) against semi-supervised learning using the same labeled examples and unlimited unlabeled examples using our learned update rule (Fig. 2, Left). We see that the learned update rule has learned to leverage unlabeled examples, achieving higher generalization accuracy. In Figure 2 (Right), the decision boundaries resulting from applying the learned update rule are nonlinear and better aligned to the underlying data distribution.

**Figure 2:**
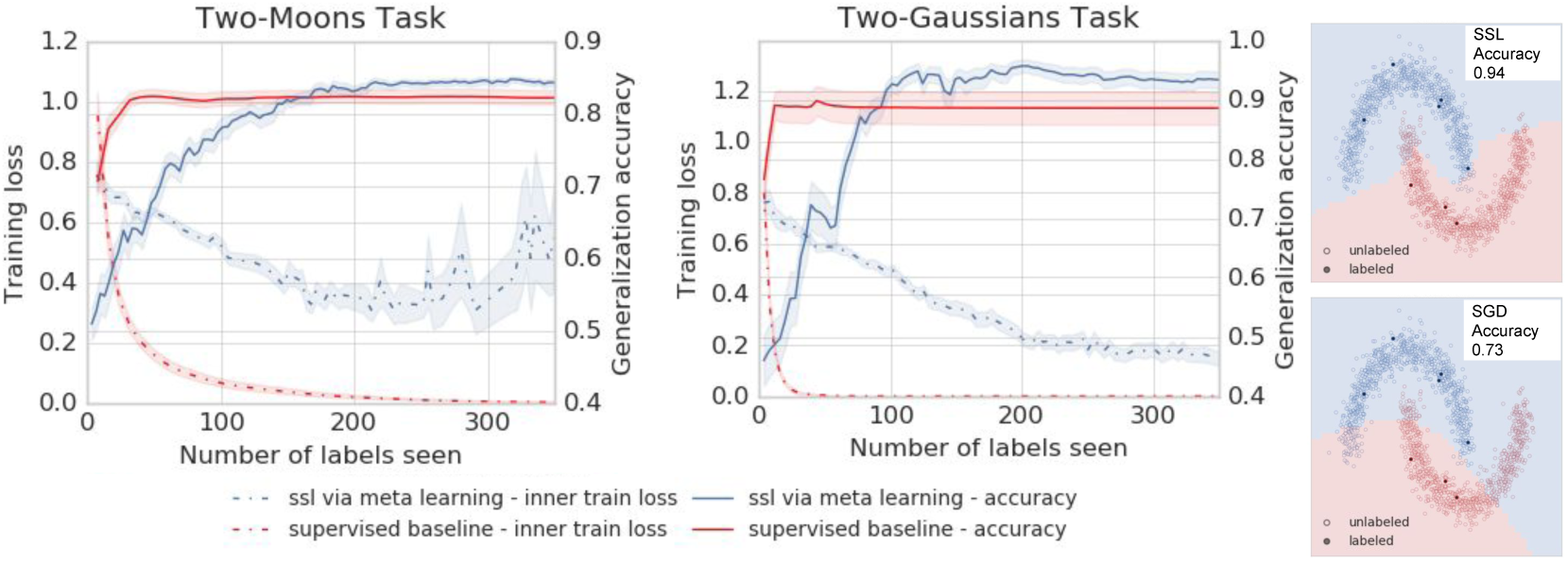
Meta-trained Semi-Supervised update rule outperforms gradient descent on toy two-moons task and generalizes to two-Gaussians. We plot the inner-accuracy and inner-training loss with respect to the number of labeled examples seen for both a supervised baseline (red) and semi-supervised learning with a learned update rule (blue), outer-trained on random two-moons tasks. The total number of labeled examples is 10, while the number of unlabeled examples available during semi-supervised learning is unlimited. Shaded regions represent standard error over 100 trials. Additionally, we show example decision boundaries on two-moons from supervised learning (bottom-right) and semi-supervised learning using a meta-learned update rule (top-right). †cosyne

We further show that the learning rule generalizes from a two moons task to random two-Gaussian tasks that it never experienced during outer-training. On this new semi-supervised learning task, it continues to outperform the supervised baseline. Note that the learned update rule solves the task in a qualitatively different way than backpropagation, and despite doing better on test accuracy performs dramatically worse on training loss.

## A Inner-Training details

Below we provide update equations describing how the inner-model weight updates are produced by the outer model:

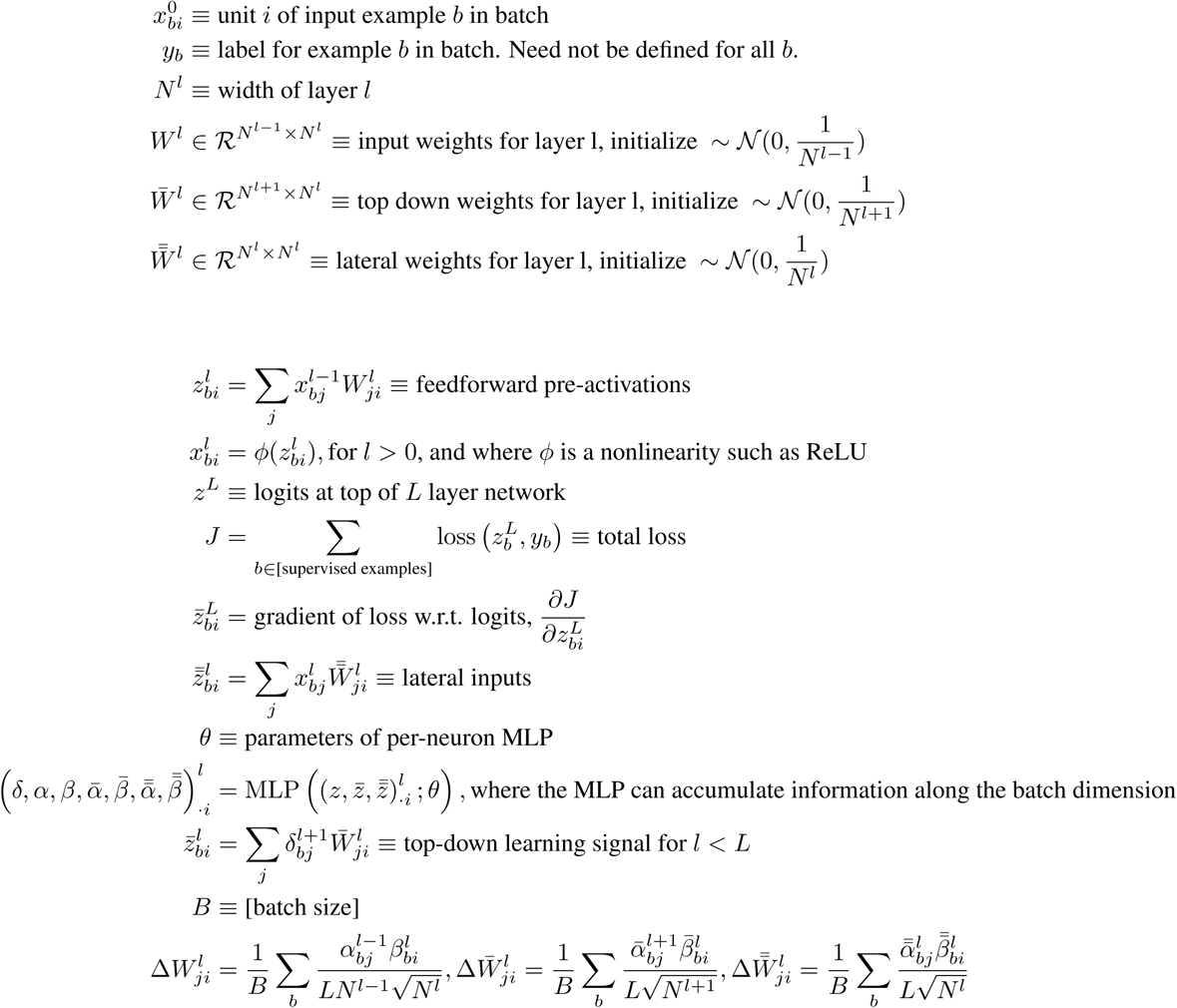

